# Bio-Aurac – an open-source browser plugin to better navigate literature content

**DOI:** 10.1101/2022.09.23.508995

**Authors:** Nick E J Etherington, Ashley J Evans, Mark P Laing, Brad Rollings, Michael J Sweeton, Alex J Whitehorn, C Southan, Gemma L Holliday, Rafael C Jimenez, Ian W Dunlop

## Abstract

**Summary:** Bio-Aurac is an open-source web browser plugin designed to support the research community in drug discovery and life sciences. The tool has been designed to help researchers, scientists, and curators to better explore, navigate and understand content from literature accessing valuable integrated information from third party resources. It identifies, highlights, and provides additionally knowledge for names of biochemical entities like genes and proteins.

**Availability and Implementation:** Bio-Aurac has been implemented using a microservice architecture which is open-source and abstracted from non-technical users by utilizing Docker containers (Nüst, 2020). It can be run with minimal prerequisites on both Chrome and Firefox browsers.

The code for installing and running the web browser plugin can be found here: https://github.com/mdcatapult/bio-aurac (Along with detailed installation instructions). A comprehensive collection of all the code involved in building this open-source project can be found: https://github.com/mdcatapult/aurac-web-plugin, https://github.com/mdcatapult/PDF-Converter, https://github.com/mdcatapult/entity-recognition

## 1 Introduction

Biological research is published at an exponential growth rate, with over 40-million articles available online at EuropePMC in July 2022. These articles are dispersed widely between various journals and preprint sites (Bethesda, 2019) with a variety of available metadata and online accessibility, from PDFs to fully open-access text. This makes gathering relevant biological information from online resources a difficult and time-consuming challenge for scientists. Most of this information, however, is accessed using a web browser which provides an opportunity for the creation of online web tools to assist in this challenging task (Pafilis, E., O’Donoghue, S., Jensen, L. et al., 2009). Named Entity Recognition (NER), or entity extraction, is a technique that aims to find specific terms or entities within unstructured text (Cho, H, 2019). In a biological context, naming convention is often challenging providing an opportunity for Machine Learning and Natural language processing (NLP) to help improve entity search efficiency (Leser and Hakenberg, 2005; Farrell, 2022). An additional challenge in biological information gathering is to then utilize the extensive online data/information repositories e.g., UniProt (The UniProt ConsortiumUniProt: the universal protein knowledgebase in 2021 -Nucleic Acids Res. 49:D1 (2021)), BioGrid (The BioGRID database: A comprehensive biomedical resource of curated protein, genetic, and chemical interactions. - Abstract - Europe PMC), *etc*. Each of these resources contain vast amounts of information that can help inform and influence a user’s understanding of the current state of knowledge, potentially leading to a better hypothesis and project definition.

Here we present an open-source resource, a web browser plugin called Bio-Aurac that identifies, highlights, and provides additionally knowledge for names of biochemical entities like genes and proteins to support researchers, scientists, and curators to better explore, navigate and understand content from literature. We also provide its required resources, including an Entity recognition/NER system that contains a single dictionary for Genes and Proteins (SwissProt) and a PDF converter. The purpose of Bio-Aurac is to be a scientist’s assistant in information gathering.

## 2 Design and implementation

Bio-Aurac is a web plugin tool that has a microservice architecture with multiple dependencies. These include a Named Entity Recognition (NER) system (and associated dictionary) and a PDF converter. Bio-Aurac has been designed with the end-user scientist in mind, thus, we have attempted to abstract as much of the technical pre-requisites as possible to streamline the process of installing and using Bio-Aurac. This abstraction allows for only a small number of prerequisites: Git and Docker, both of which are freely available with detailed installation instructions available. See GitLab repository (https://github.com/mdcatapult/bio-aurac) for full installation instructions. Once installed, the plugin can then be added to a web browser (Chrome is recommended) and a user can start using Bio-Aurac.

Once installed, the user simply needs to visit a website and open the pop-up window (Supplementary Figure 1). If there are terms in the text that are Gene and Protein entities that exist within the Swiss-Prot dataset and selected species of interest (*Homo sapiens* is the default), Bio-Aurac will identify them using the supplied dictionaries and highlight them (Supplementary Figure 2). Clicking on a highlighted entity will open and then populate the Bio-Aurac sidebar (Supplementary Figure 3) which contains more detailed information on the entity, including known synonyms, sequence information, general data, and a pre-selected list of websites against which the entity of interest can be queried (Supplementary Figures 4-7). The full list of resources to which Bio-Aurac links out too can be found in the Supplementary Table 1.

Many manuscripts, especially pre-prints and older papers are available only as PDFs. Thus, we have designed Bio-Aurac with a PDF converter to allow these valuable resources to also be marked up. In the popup, the PDF conversion section can be expanded and then a URL to the PDF of interest can be entered into the input field to be converted (Supplementary Figure 8). After the conversion is complete a new browser tab will be opened containing the PDF contents that can then be highlighted using Bio-Aurac. (Supplementary Figure 9-10).

Another key feature of Bio-Aurac is the ability to filter highlighted results on a species of interest. *Homo sapiens* is set by default but if the user works using a different animal model, then there are eight to choose from (Supplementary Table 2). When a user switches configuration to a species of interest, highlights the page, and then selects an entity of interest, all the information and links contained in the sidebar card will be specific for the species selected (Supplementary Figures 11-13).

Bio-Aurac is implemented using three core individual components:

### Web-browser plugin

The web-browser plugin is built using Typescript (https://www.typescriptlang.org/) and Angular (https://angular.io/) with four primary components: the popup, the sidebar, the background, and the content script. The plugin can be added to both Chrome and Firefox browsers. The plug-in code is available from https://github.com/mdcatapult/aurac-web-plugin

### Entity recognition system

The entity recognition system is responsible for scanning the webpage text and distinguishing between entities of interest and dis-interest using NER under the hood. It is a web-service written in Golang (https://go.dev/) and made up of 3 individual components: the Recognition API (Application Programming Interface), the Dictionary Recognizer, and the Dictionary Importer. When entity recognition is initialized, the Dictionary Importer will add all the Swiss-Prot database entries into a Redis (https://redis.io/) dictionary instance. Then the recognition API and Dictionary Recognizer will initialize, allowing requests to be sent from the web-browser plugin to the recognition API. Streams of tokens are then sent via a gRPC (https://grpc.io/) method to the Dictionary Recognizer which in turn receives these token streams and checks for presence within the Redis dictionary. If an entity is found this is sent back from the recognizer to the API and then back to the web-browser plugin via an http response which then allows the web-plugin to highlight the entities on the web page. The code for this component is available from https://github.com/mdcatapult/entity-recognition.

### PDF Converter

The PDF converter is a dependency of the web-browser plugin. Requests are sent to the PDF converter when a URL is entered into the plugin’s popup PDF conversion input. When the request is received, the PDF is converted to HTML and then populated into a new browser tab. Bio-Aurac can then highlight the HTML version of the PDF. The PDF converter is a web service that is built upon the PDF.js open-source project (https://github.com/mozilla/pdf.js) and is available from (https://github.com/mdcatapult/PDF-Converter).

## 3 Tools and future directions

Bio-Aurac is an open-source web-browser plugin that can be installed and set up within minutes. By abstracting the core requirements into an easy-to-use architecture, a user does not need to be a computer scientist to get this tool up and running.

We have demonstrated its use based on Swiss-Prot gene and protein names for eight core species. By making the code available, the underlying entities that can be recognized may be expanded to new species, new entity types, or larger datasets.

After providing a foundation we now encourage the opensource community to expand and personalise this tool further.

All these individual tools have been made available to the community through GitHub and are all freely available and released under the Apache 2.0 license.

## Acknowledgements

We would like to thank our colleagues at MDC (Medicines Discovery Catapult) especially Chris Southan, Emily Offer, Gayle Marshall, Sophie Nyberg, Emma Jones, Priya Viswanathan and Lucy Frost for testing and providing feedback regarding Bio-Aurac, Rob Bevan for NER benchmarking work and John Overington for initial discussion.

