## Supplementary Figures for "Bio-Aurac – an open-source browser plugin to better navigate literature content"

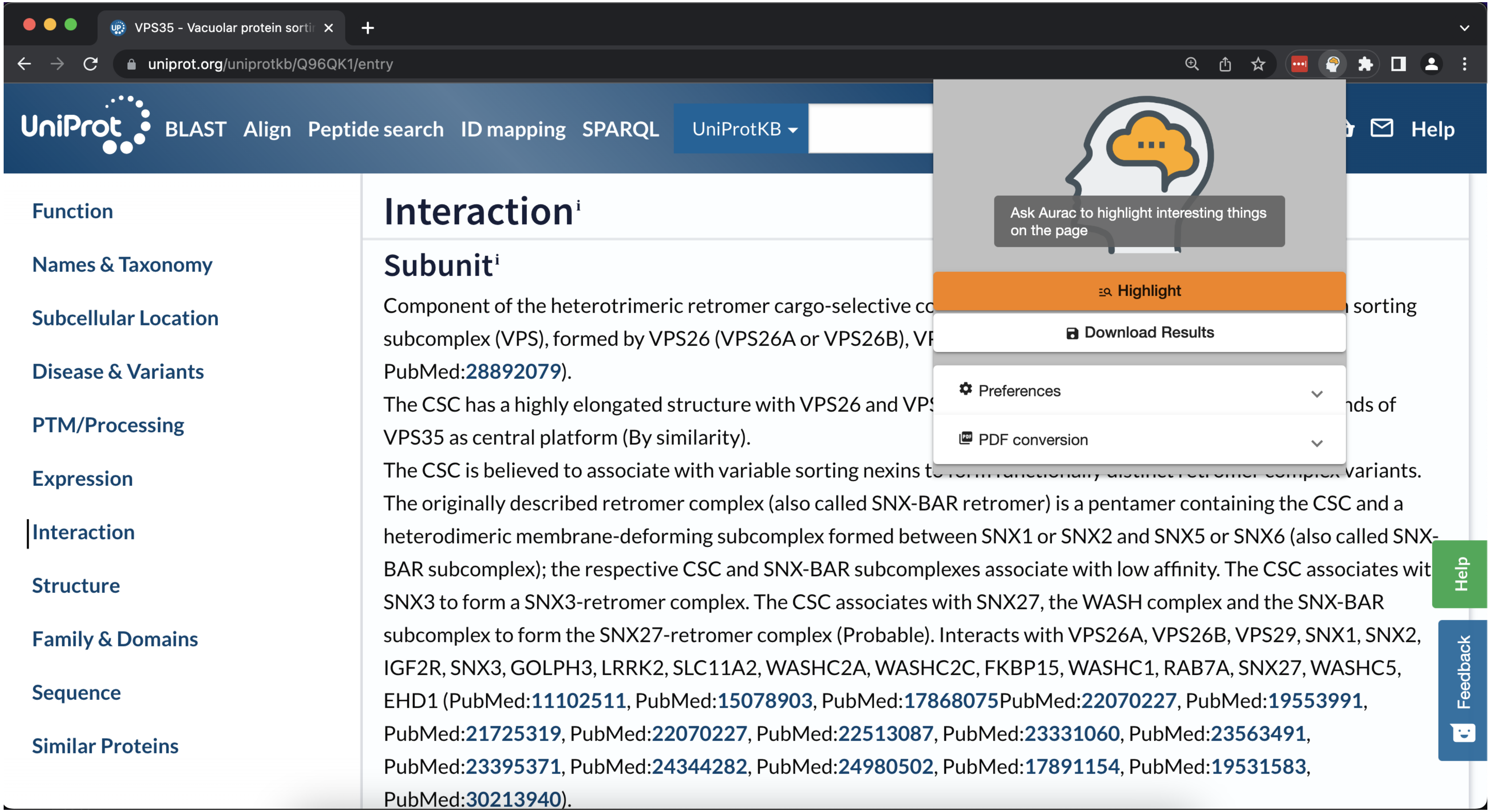


Supplementary Figure 1 – An example of a user visiting a uniport webpage and then clicking on the Bio-Aurac plugin

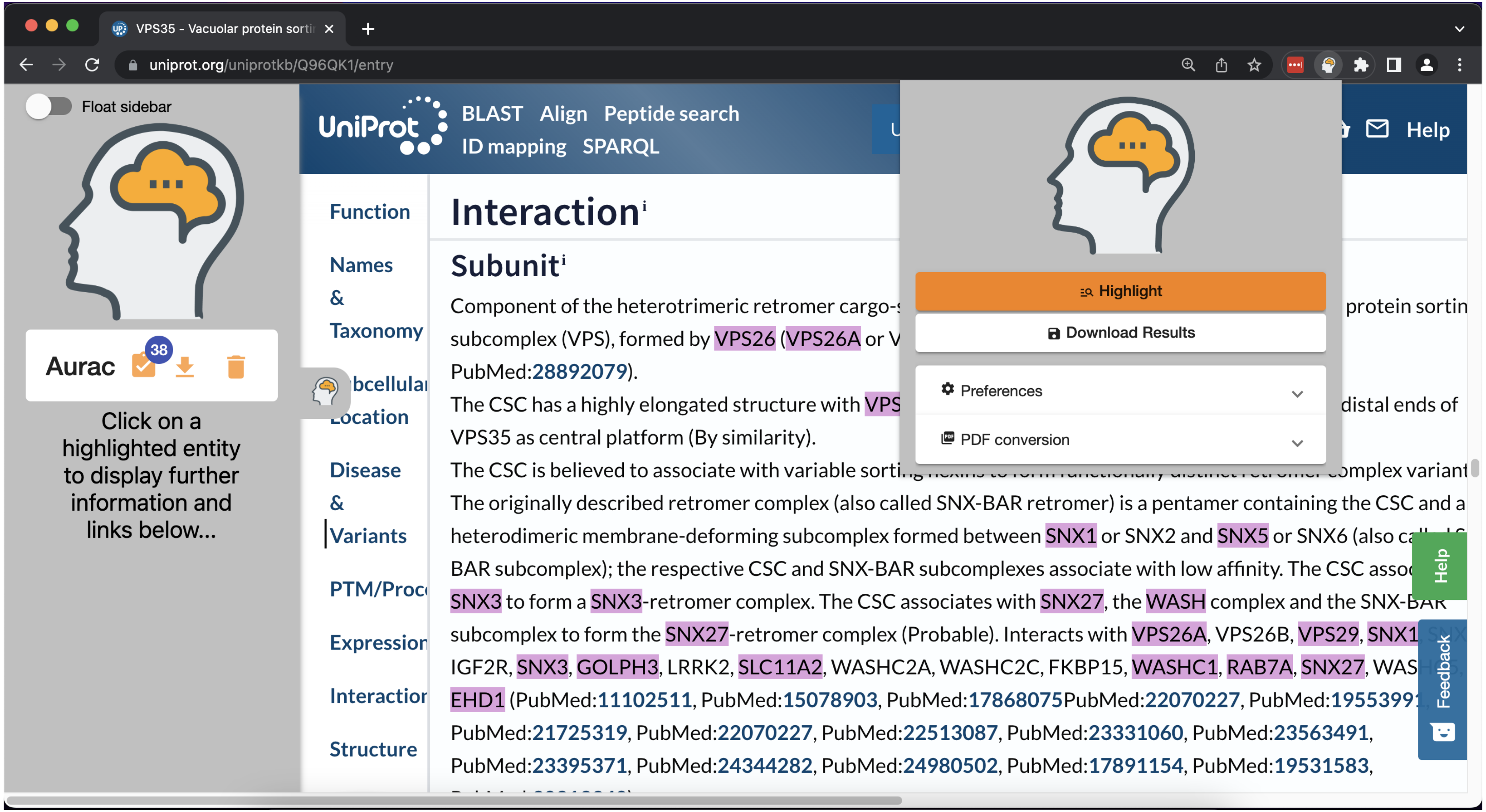


Supplementary Figure 2 – A uniport webpage after protein/gene highlighting by Bio-Aurac.

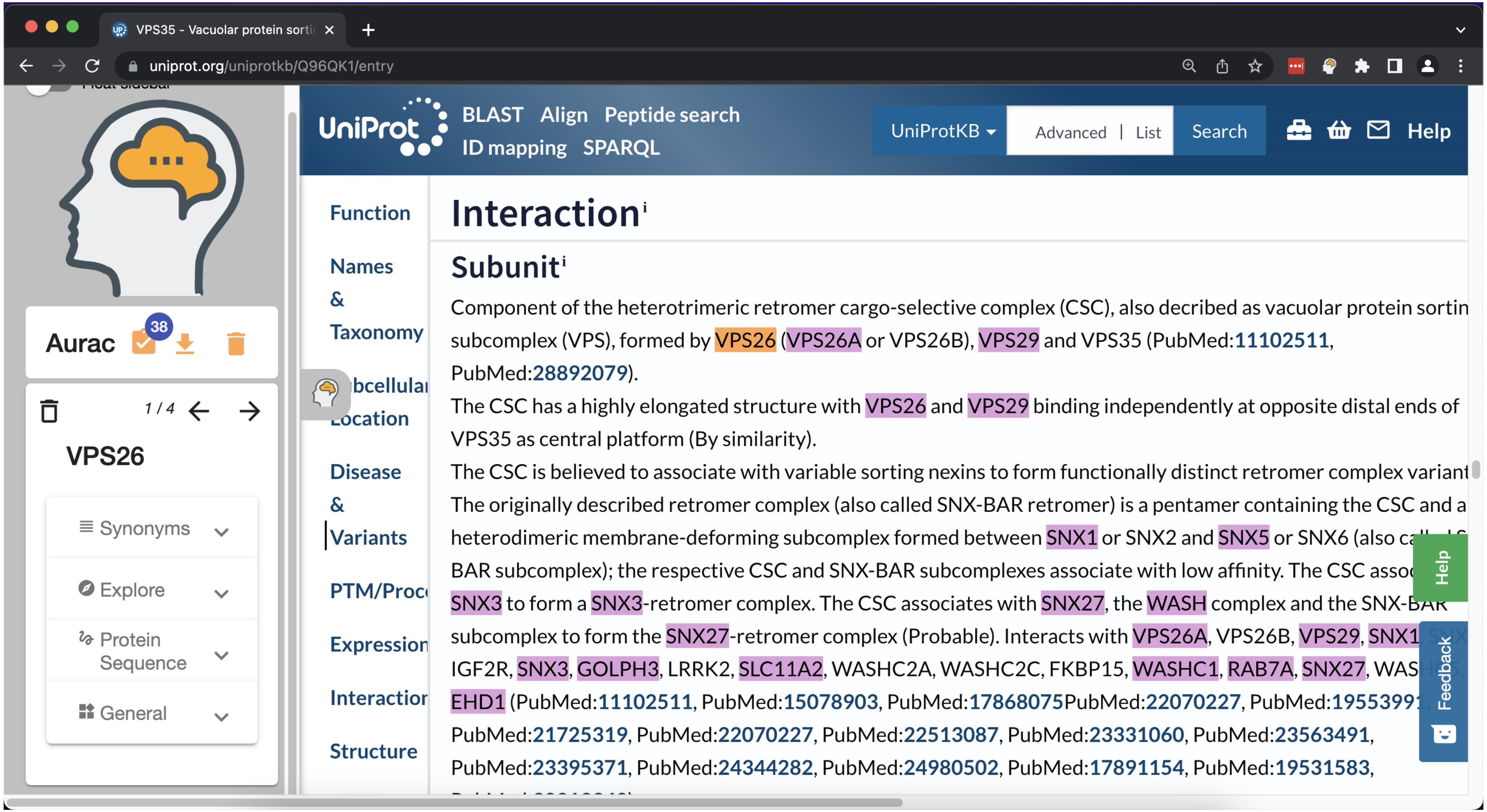


Supplementary Figure 3 – The sidebar is populated with the entity 'VPS26' when a user selects if from the main page

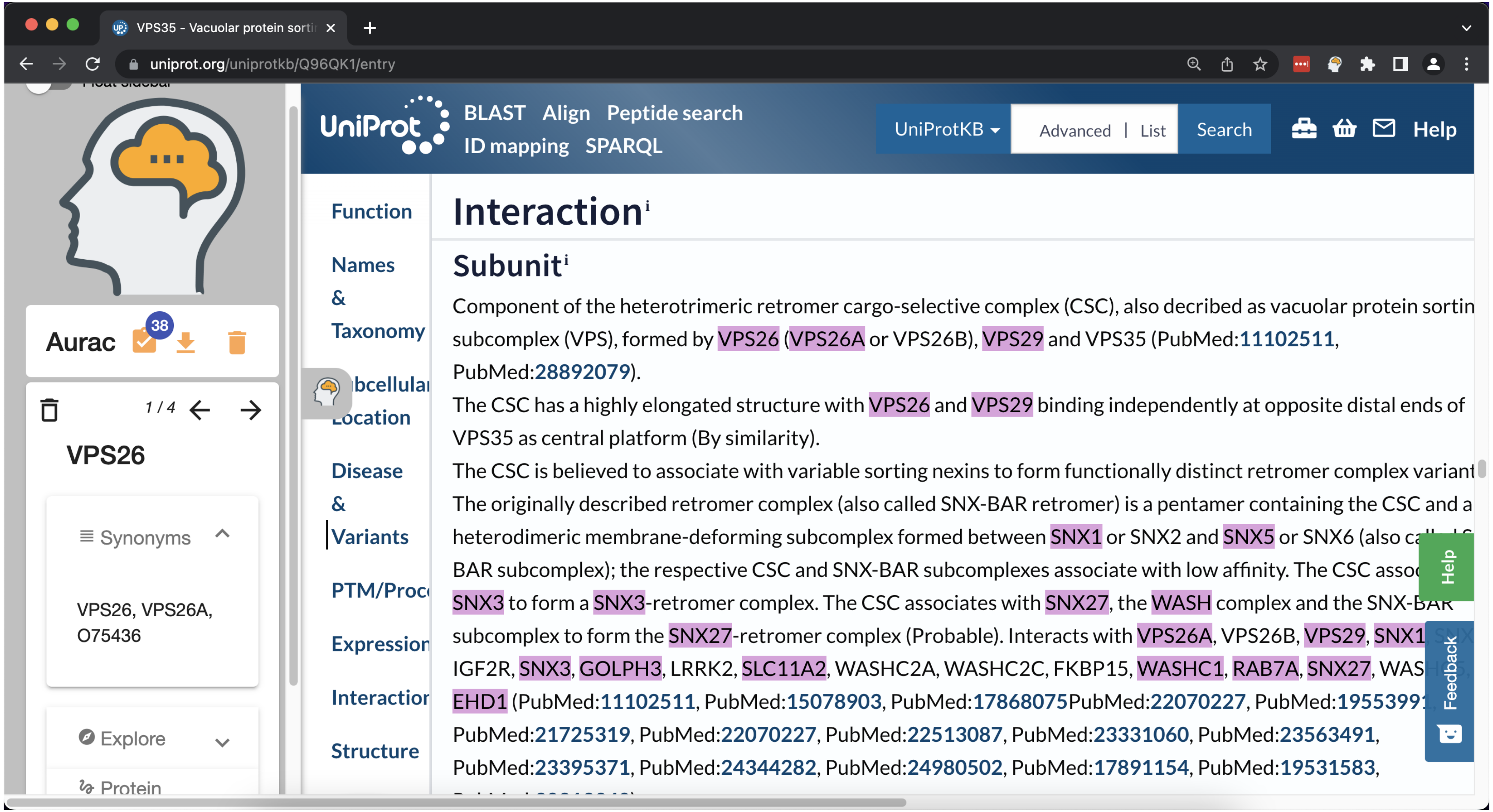


Supplementary Figure 4 – A user exploring the different synonyms that appear on the current page for 'VPS26'

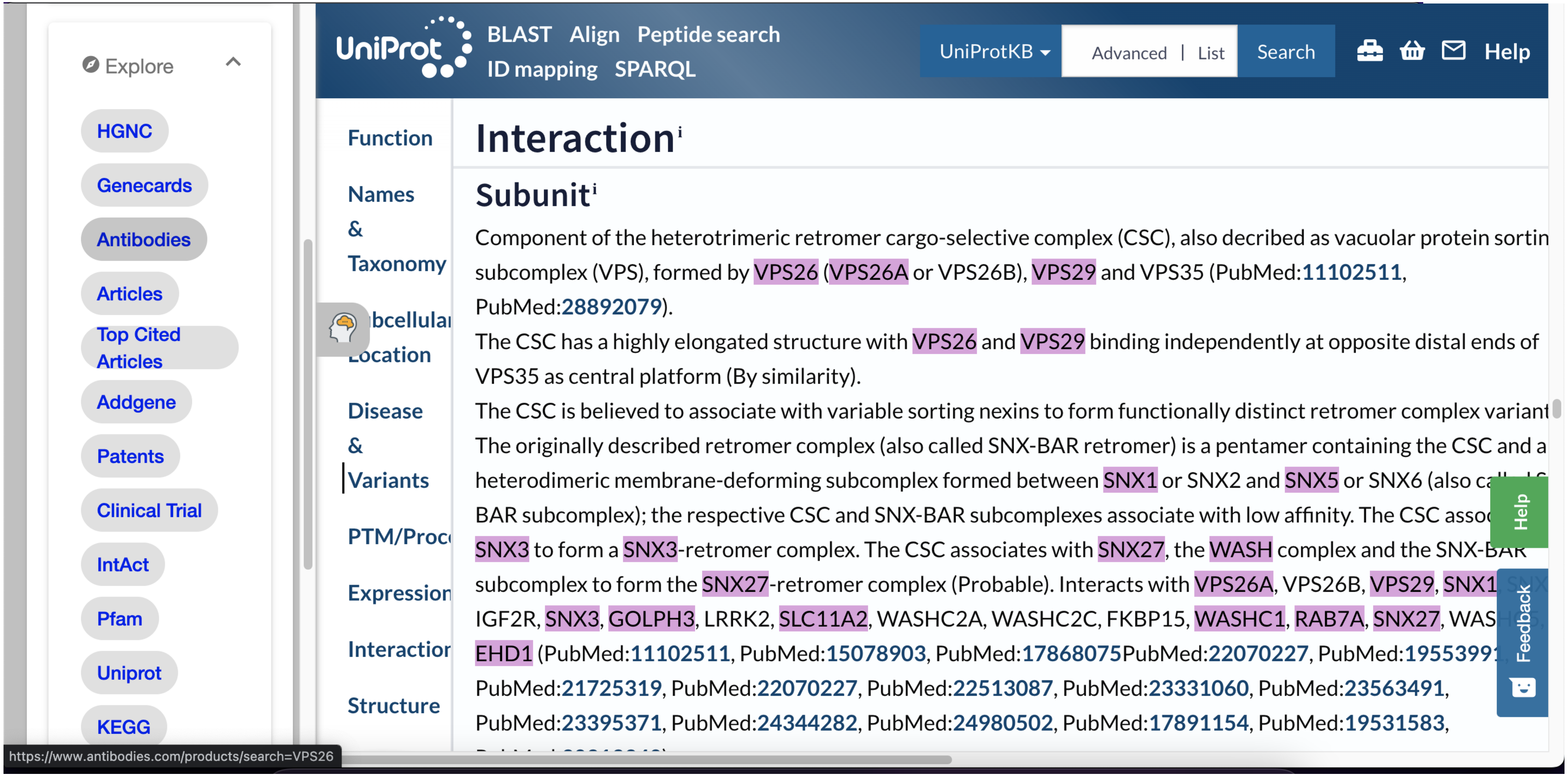


Supplementary Figure 5 – A user exploring the range of available links out too biological resources.

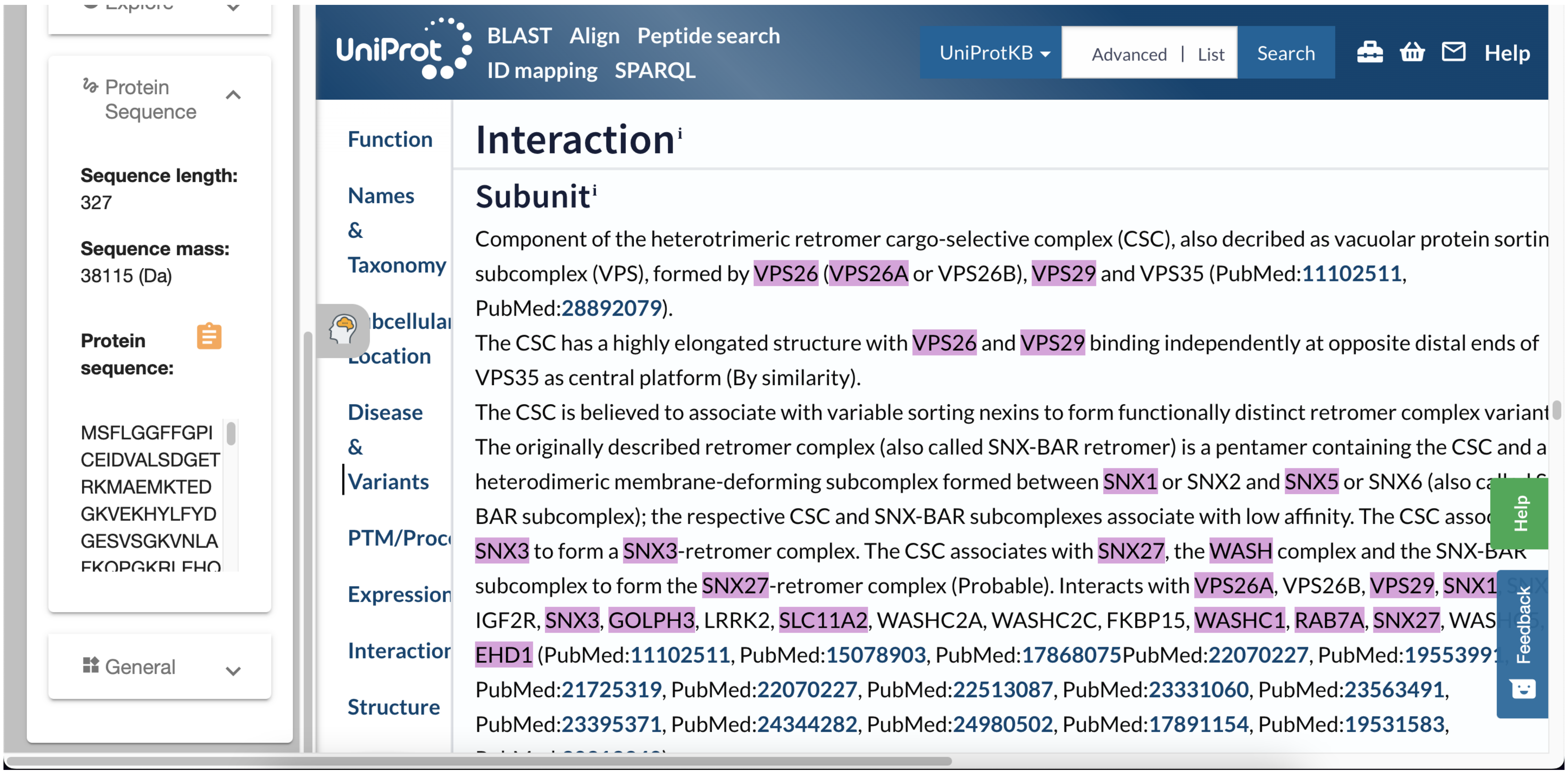


Supplementary Figure 6 – A user discovering the protein sequence length, mass and sequence of 'VPS26'.

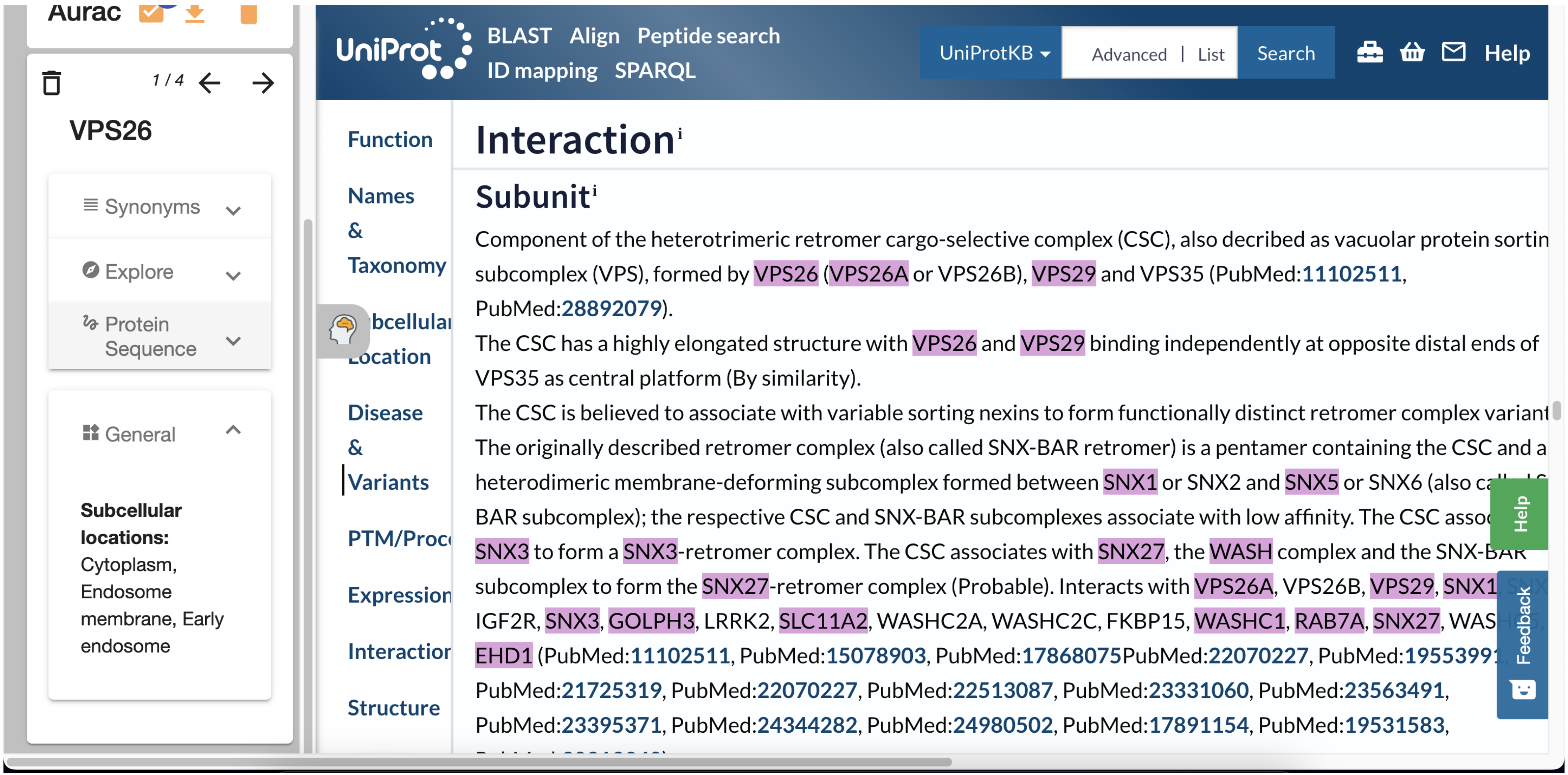


Supplementary Figure 7 – A user investigating general information about ‘VPS26’ including the subcellular locations.

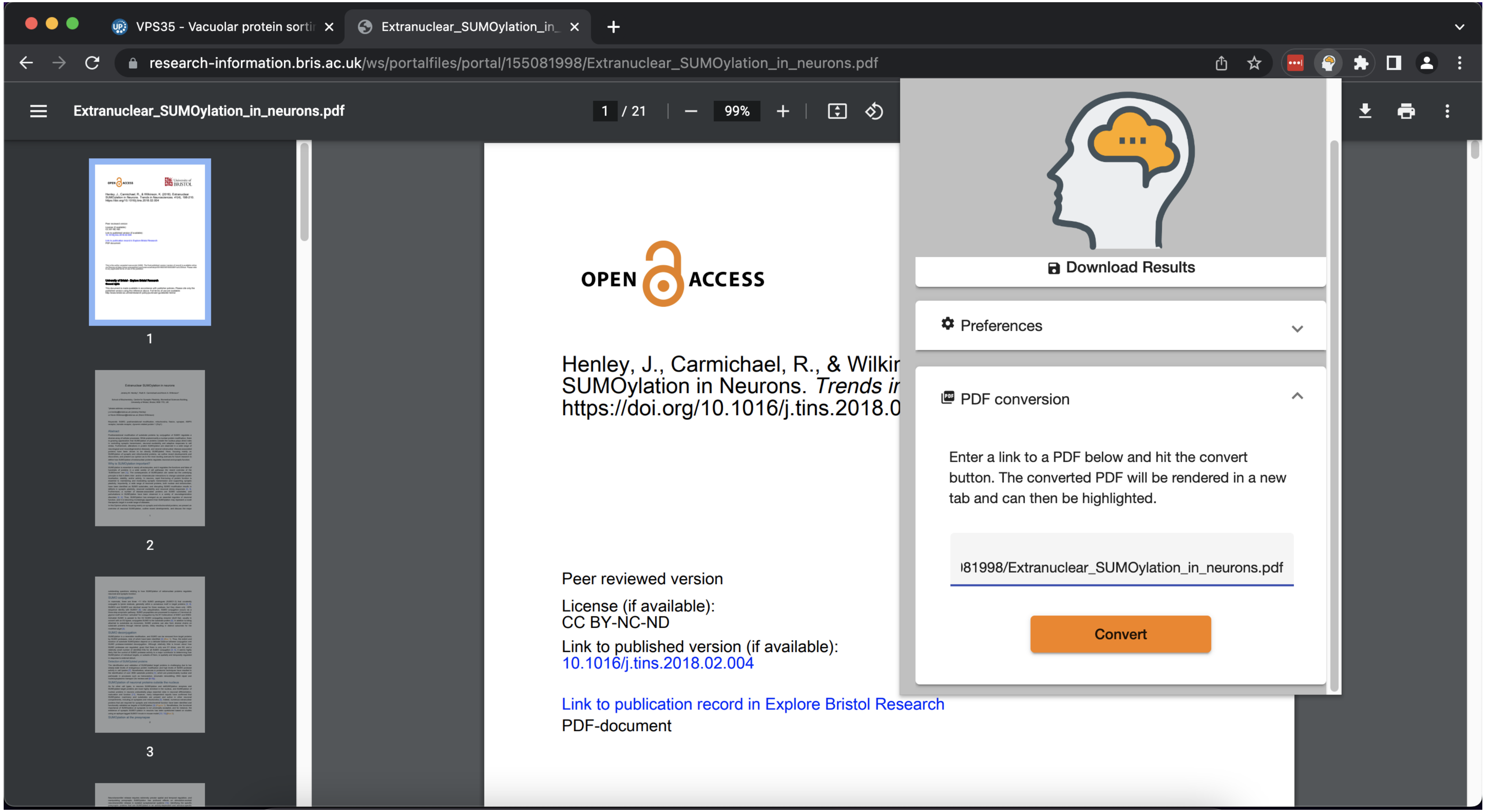


Supplementary Figure 8 – A user entering a PDF URL into the PDF converter field in the Bio-Aurac popup.

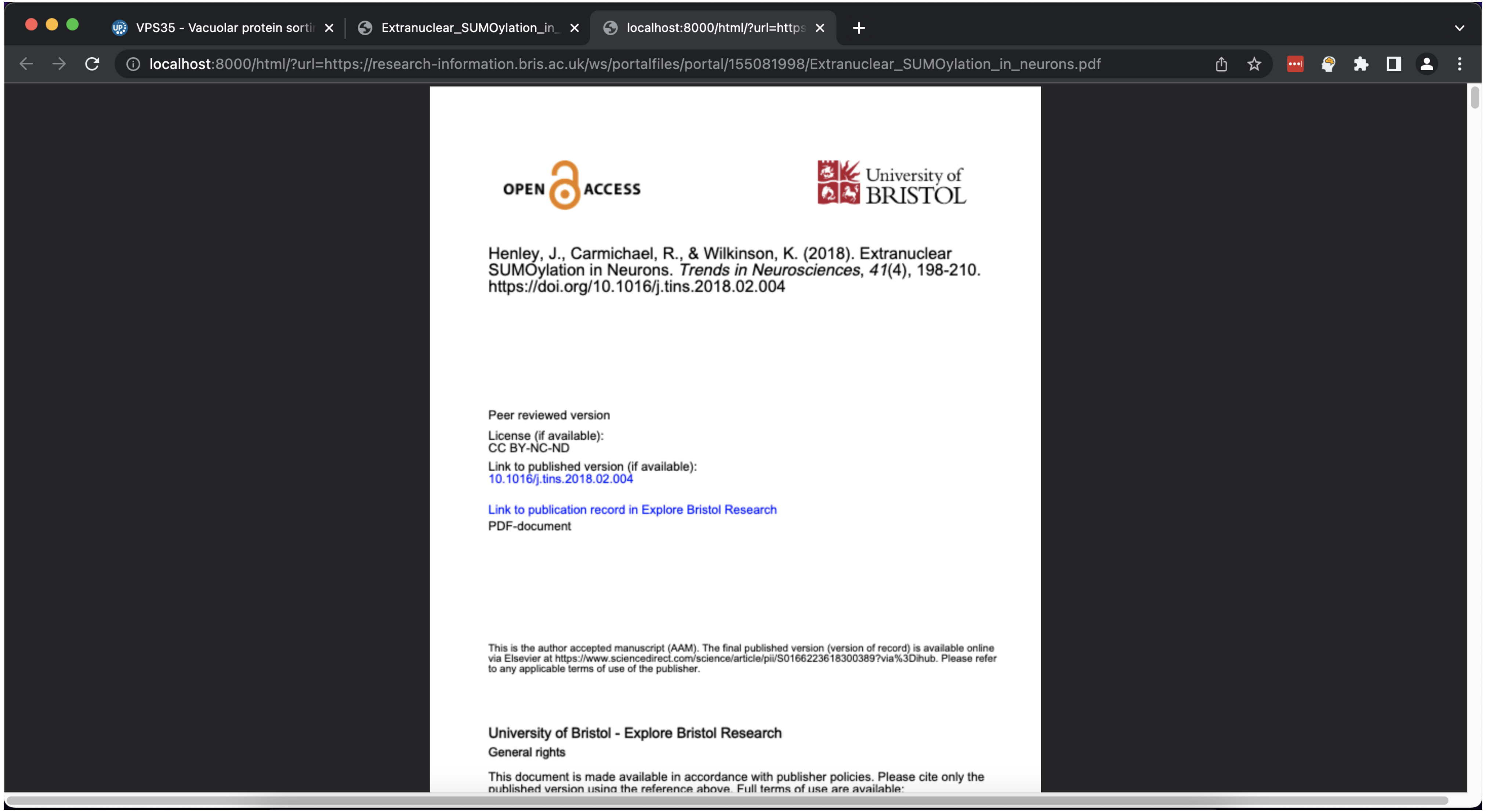


Supplementary Figure 9 – After pressing convert a new tab appears with the content of the converted PDF.

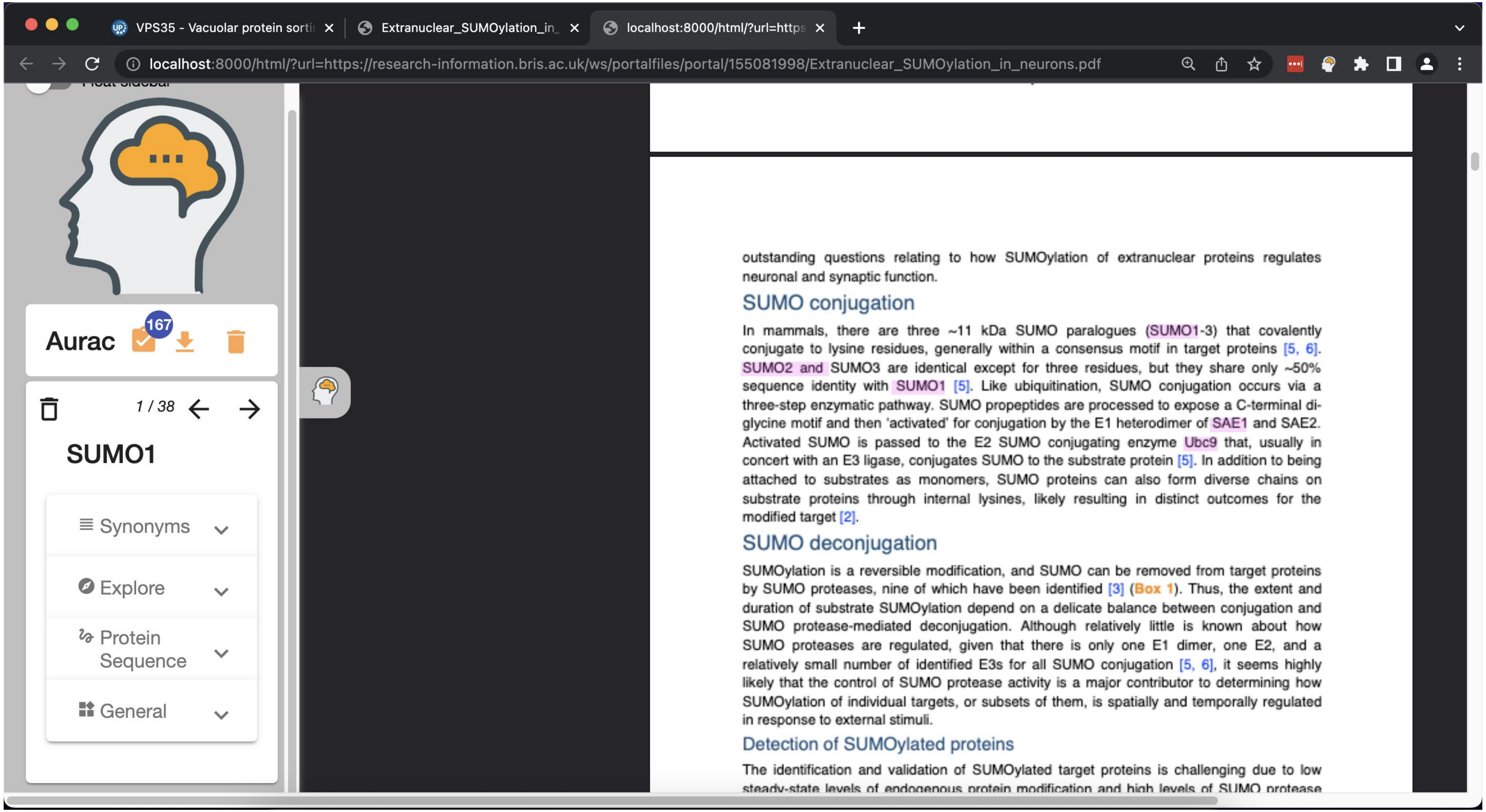


Supplementary Figure 10 – A user can now use the Bio-Aurac plugin to highlight the converted PDF.

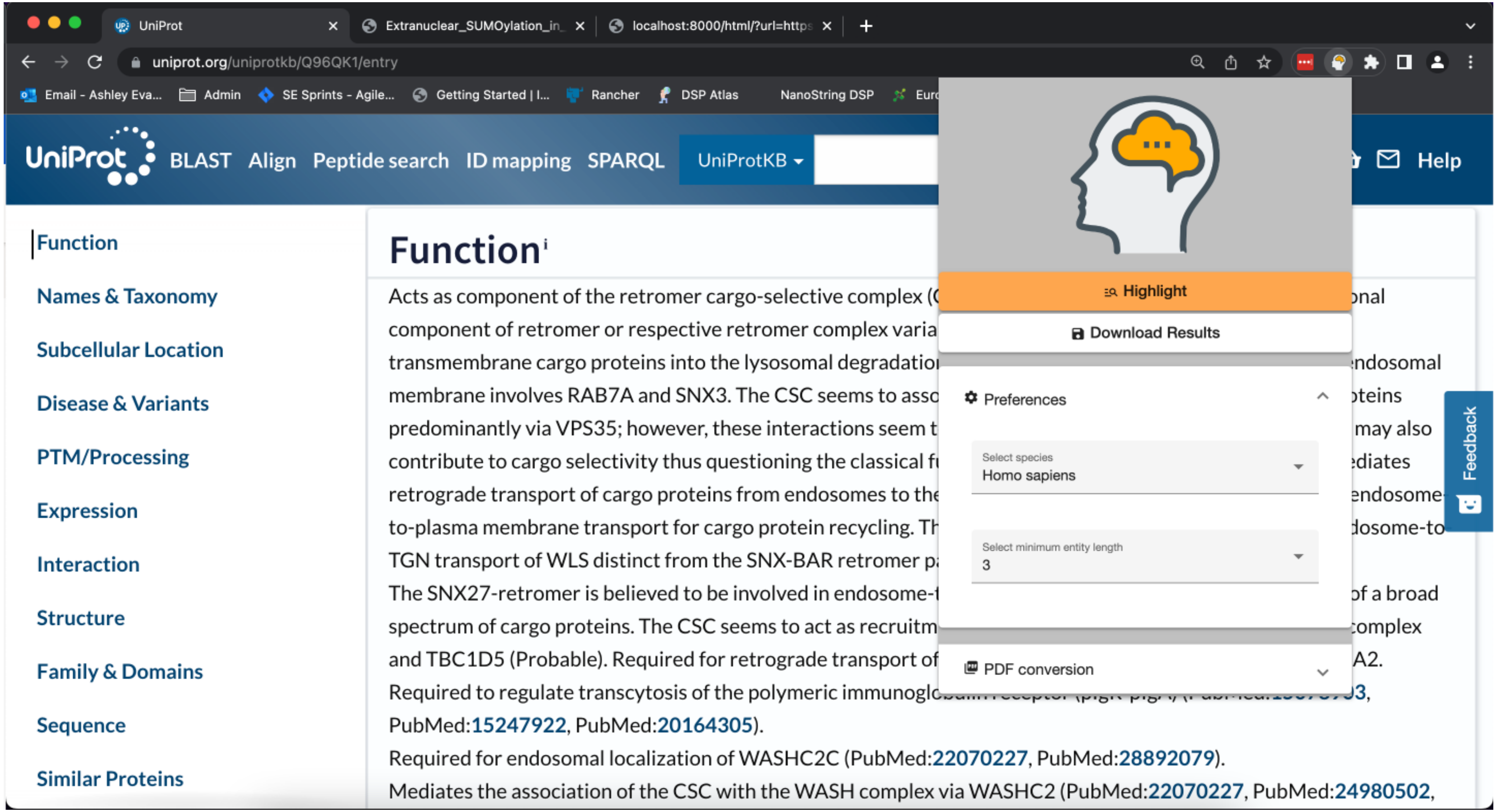


Supplementary Figure 11 – A user can change both the species they are interested in searching for entities and the minimum entity length of each entity.

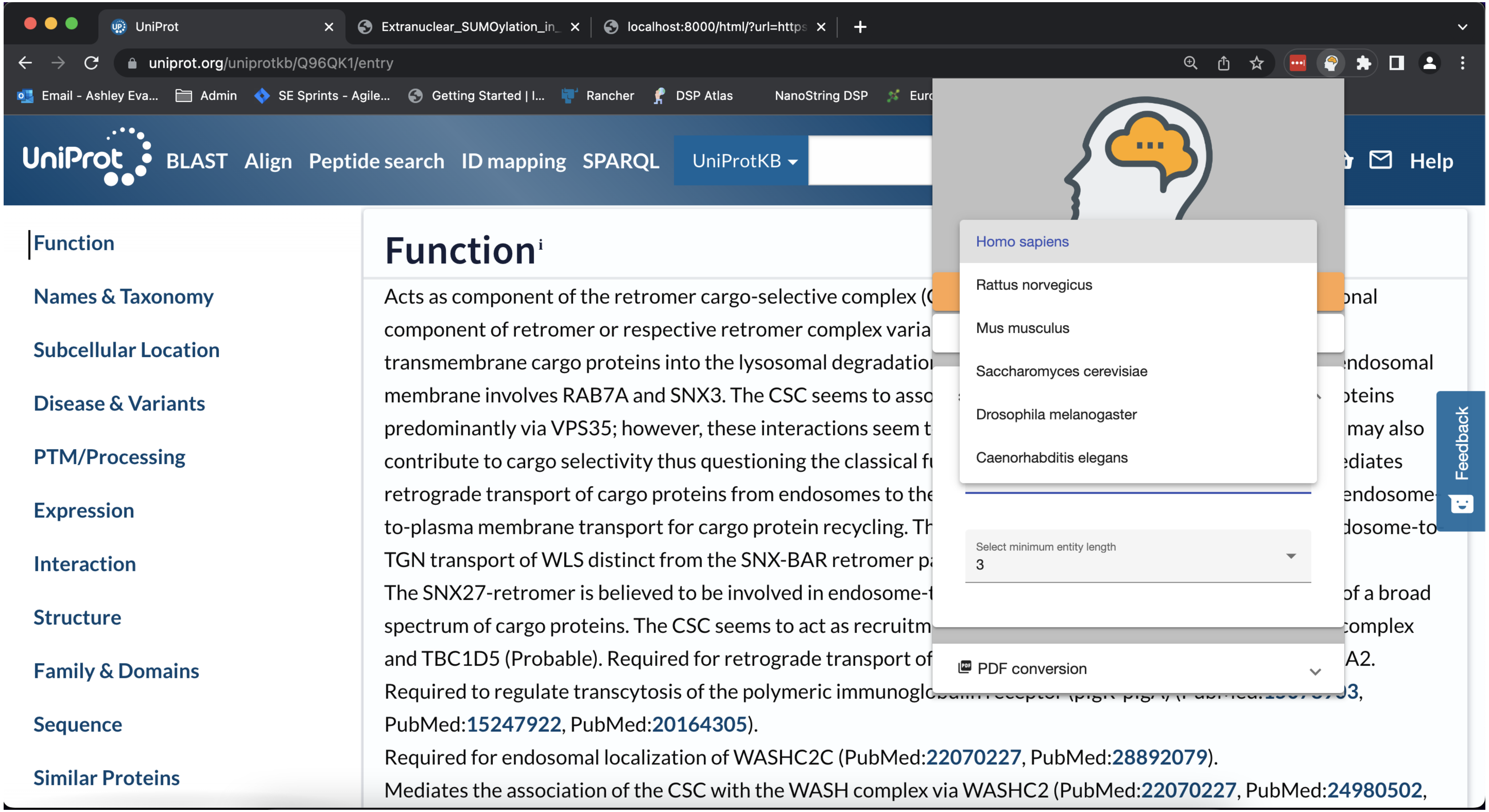


Supplementary Figure 12 – There are 8 species that can be selected by the user depending on their research interests.

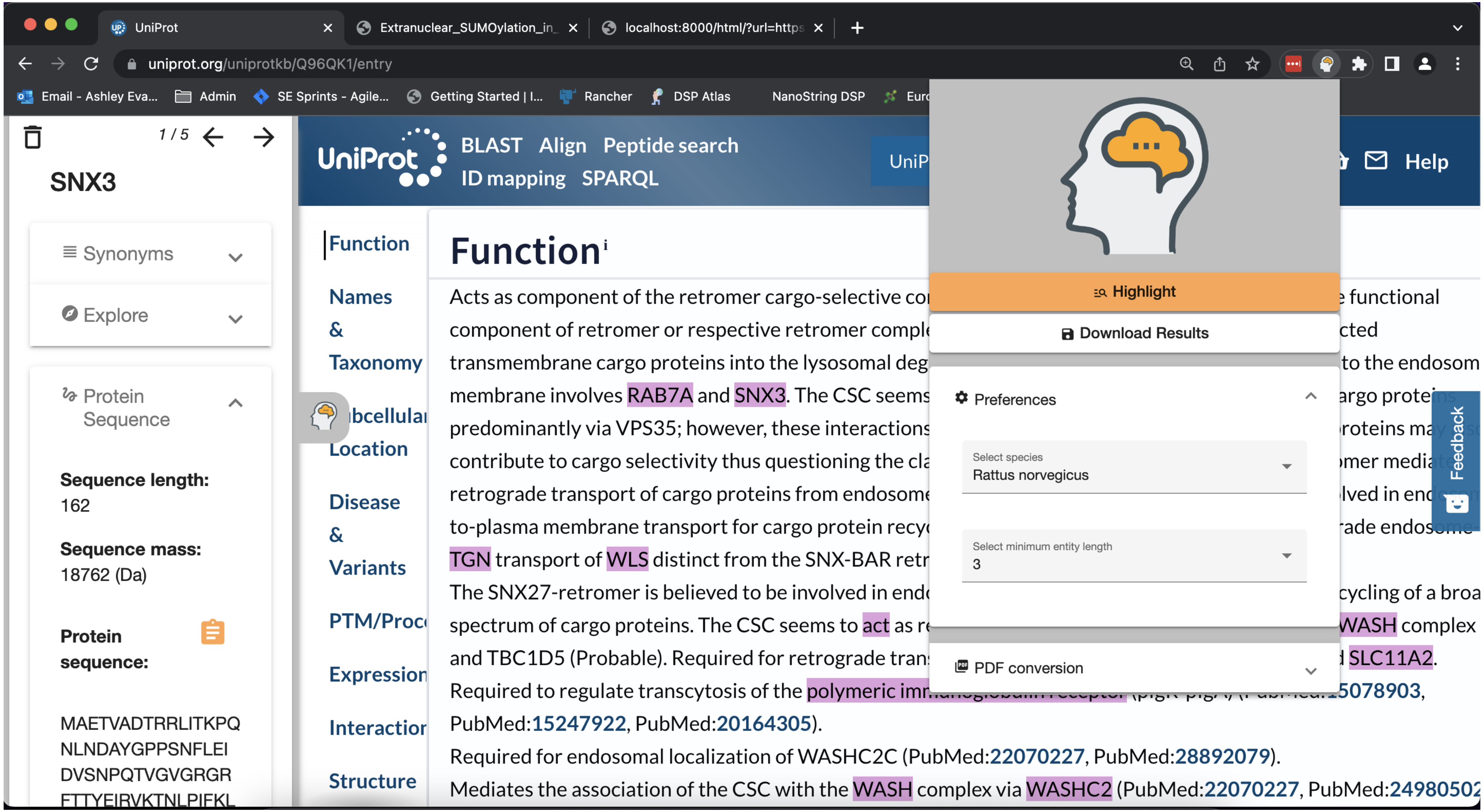


Supplementary Figure 13 – A user has selected Rattus norvegicus and the page has re-highlighted with entities that the database has Rattus norvegicus specific information.

| Label | Url (Example using RAB7A as the selected entity) |
|---|---|
| **HGNC** | https://www.genenames.org/data/gene-symbol-report/#!/hgnc_id/RAB7A |
| **Genecards** | https://www.genecards.org/cgi-bin/carddisp.pl?gene=RAB7A |
| **Antibodies** | https://www.antibodies.com/products/search=rab7a |
| **Articles** | https://pubmed.ncbi.nlm.nih.gov/?term=RAB7A&sort=date |
| **Top Cited Articles** | https://app.dimensions.ai/discover/publication?search_mode=content&search_text=RAB7A&search_type=kws&search_field=text_search&order=times_cited |
| **Addgene** | https://www.addgene.org/search/catalog/plasmids/?q=RAB7A |
| **Patents** | https://patents.google.com/?q=RAB7A |
| **Clinical Trial** | https://clinicaltrials.gov/ct2/results?cond=&term=RAB7A&cntry=&state=&city=&dist= |
| **IntAct** | https://www.ebi.ac.uk/intact/search?query=P09527 |
| **Pfam** | http://pfam.xfam.org/family/PF00071 |
| **Uniprot** | https://www.uniprot.org/uniprotkb/P09527/entry |
| **KEGG** | https://www.genome.jp/entry/rno:29448 |
| **NCBI** | https://www.ncbi.nlm.nih.gov/gene/29448 |
| **BioGRID** | https://thebiogrid.org/248093 |
| **Ensembl** | https://www.ensembl.org/Rattus_norvegicus/Transcript/Summary?g=ENSRNOG00000012247;r=4:120461963-120506850;t=ENSRNOT00000016432 |
| **GeneTree** | https://www.ensembl.org/Multi/GeneTree/Image?gt=ENSGT00940000155864 |

Supplementary Table 1 – A list of resources a user can link out too from Bio-Aurac

| **Supported Species** |
|---|
| Homo sapiens |
| Rattus norvegicus |
| Mus musculus |
| Saccharomyces cerevisiae |
| Drosophila melanogaster |
| Caenorhabditis elegans |
| Xenopus tropicalis |
| Danio rerio |

Supplementary Table 2 – A list of supported species in Bio-Aurac
